# Localized oxygen control in a microfluidic osteochondral interface model recapitulates bone-cartilage crosstalk during osteoarthritis

**DOI:** 10.1101/2023.11.20.567822

**Authors:** Louis Jun Ye Ong, Antonia Rujia Sun, Zhongzheng Wang, Jayden Lee, Indira Pradasadam, Yi-Chin Toh

## Abstract

Osteoarthritis (OA) is characterized by the dysregulation of the osteochondral interface between bone and cartilage. *In vitro* models that accurately mimic this interface hold great potential for understanding OA pathophysiology and screening therapeutic agents. Presently, research efforts have focused on emulating heterogeneity in structural and mechanical attributes of the extracellular matrix (ECM) at the osteochondral interface. However, the precise simulation of differential oxygen gradients experienced by chondrocytes and osteoblasts in vivo remains a substantial obstacle for modeling osteo-chondral interactions effectively. To overcome this limitation, we show that micropatterned granular hydrogels, which are small microgel particles swelled in liquid culture media to create a shear-yielding jammed-packed solid, can be used to control the delivery of oxygen scavenging agents in a simple and scalable manner. Hypoxic granular hydrogels formulated with Oxyrase™ could maintain <1% oxygen concentration in a conventional cell culture incubator. Primary human chondrocytes maintained in the hypoxic hydrogels expressed a more anabolic phenotype similar to those cultured in a hypoxic incubator. The granular hydrogels can be readily patterned in a microfluidic device to generate a localized hypoxic environment, mimicking the differential oxygen levels at the osteochondral tissue interface (i.e. osteoblast at 20% and chondrocyte at 2% oxygen). Using this microfluidic coculture model, we paired healthy human chondrocytes with osteoblasts isolated from non-sclerotic and sclerotic subchondral bone to investigate how oxygen environment modulates osteoblast-chondrocyte crosstalk during OA. In a differential oxygen environment, the osteoblast-chondrocyte co-culture model showed sclerotic osteoblasts inducing chondrocyte collagen expression changes through increased MMP13 and ADAM15 production, unlike in a uniform normoxic oxygen environment, where the change was driven by altered collagen gene expression favoring Type I over Type II collagen. Furthermore, differential oxygen conditions enabled the identification of extensive transcriptional alterations induced by sclerotic osteoblasts, which involved inflammatory NF-κβ, TGF-β/BMP, and IGF signaling pathways, that was otherwise not detectable in a uniform normoxic co-culture. The microfluidic model with localized oxygen variations effectively mimics physiologically relevant osteoblast-chondrocyte crosstalk, providing valuable insights into OA pathophysiology.

## Introduction

The osteochondral interface between bone and cartilage in the joint is one of the most ubiquitous tissue interfaces in the human body. It has clinical importance because many joint diseases such as osteoarthritis (OA) involve dysregulation of bone-cartilage crosstalk across this tissue interface [1]. In addition, implanted acellular or tissue grafts would need to integrate with the native osteochondral interface tissue [2]. Therefore, there have been extensive efforts to engineer the osteochondral interface by establishing osteocyte-chondrocyte co-cultures in extracellular matrices (ECM) with localized control of matrix stiffness, porosity, topography, and biochemical cues [2]. However, it has not been possible to recapitulate local variation in oxygen concentrations across the osteochondral tissue interface even though it is well-established that chondrocytes are exposed to 1-2% oxygen (hypoxic) concentration while neighboring osteoblasts are exposed around 9% oxygen levels *in vivo* [3]. Maintenance of these physiological oxygen concentrations is important for both cultured chondrocytes and osteoblasts to maintain their tissue-specific phenotype and functions *in vitro*. For instance, chondrocyte anabolism, which is important for forming hyaline cartilage, is enhanced under a hypoxic environment, and inhibited at normoxia [4, 5]. In contrast, bone tissue showed inhibited bone mineralization and regeneration when exposed to a low oxygen concentration [6]. Therefore, there is a need to locally control oxygen concentrations in engineered osteochondral tissue models to recapitulate physiologically relevant crosstalk between bone and cartilage during normal or diseased conditions.

The conventional method of controlling oxygen concentrations in cell culture incubators) and have one pre-set concentration (e.g., 5% for hypoxic and 20% for normoxic conditions), and thus cannot mimic differences in oxygen levels across the osteochondral interface. Microfluidic devices have been developed to generate oxygen concentration gradients in cell cultures by designing channel networks to directly flow gas [7] or chemical reducing agents [8], which are separated from the culture medium by a gas permeable membrane. Equilibration between the gas or chemical reducing agent with the culture medium results in a reduction of dissolved oxygen in the culture media. For example, Ingber’s team relied on a gas network containing inert nitrogen gas to establish a co-culture of gut epithelium maintained under hypoxia with vascular endothelium maintained under normoxia [7]. Chen et al. perfused a biocompatible oxygen scavenger solution to first treat culture media to generate an oxygen gradient before exposure to cell culture within the device [8]. However, both approaches require a constant flow of nitrogen gas or oxygen scavenging solution, which necessitates the use of enclosed microfluidic devices, tubing and pumps. Such perfusion setups increase the engineering complexity of the *in vitro* tissue constructs, which limit their practical deployment and scale-up in routine biological labs.

To overcome this limitation, we posit that the use of patternable solids, such as hydrogels, can offer a simple and scalable means to spatially control the delivery of oxygen scavenging agents, thereby generating localized hypoxic environments to recapitulate the differential oxygen environments at the osteochondral tissue interface. To this end, we employ a class of granular hydrogel, whereby small (∼20 μm diameter) polyacrylate microgel particles are swelled in liquid culture media to create a jammed-packed solid [9]. This hydrogel does not require any crosslinking since it is held together by reversible hydrogen bonds [10]. Hence, there is no risk of damaging the oxygen scavenger or other soluble factors in the culture medium. In addition, this granular hydrogel exhibits shear-yielding properties rendering it amenable to being patterned within a microfluidic device or by bioprinting [10]. Here, we demonstrate the feasibility of patterning oxygen-scavenger-laden granular hydrogels (hereafter referred to as hypoxic hydrogels) in a microfluidic device to recapitulate differential hypoxic and normoxic conditions present at the osteochondral interface. More importantly, we show that the microfluidic osteochondral co-culture model maintained under differential oxygen levels could recapitulate physiologically relevant bone-cartilage crosstalk in the context of OA pathophysiology.

## Materials and Methods

### Reagents

Unless otherwise stated, all chemicals and reagents were purchased from Sigma-Aldrich Pte Ltd, Australia. Polyacrylate microgel (trade name Carbopol 940) is procured via Lubrizol, United States.

### Granular hydrogel formulation

2.0 w.t./v % granular hydrogel was first prepared by weighing out the required amount of Carbopol 940 polyacrylate microgel powder. The powder was sterilized with UV exposure for 60 mins. The UV-treated microgel powder was then dissolved in deionized water under gentle stirring. 1M sodium hydroxide was added dropwise to neutralize the acidic polyacrylate microgel solution until pH 7, at which the solution will swell into a hydrogel. The hydrogel was centrifuged at 1000 ×g for 10 minutes to remove any trapped air bubbles.

### Normoxic and hypoxic granular hydrogel formulation

To prepare 1% (wt/v %) normoxic media hydrogels, we mixed 1-part 2% (wt/v %) hydrogel, which was swelled and neutralized in DI water, with 1-part 2× low glucose DMEM culture medium (Thermo Fisher Scientific, Australia). Hypoxic hydrogels were prepared in similar way except sodium sulfite or Oxyrase^TM^ was first added to the DMEM culture medium. This strategy was adopted to minimize the influence of pH fluctuations in the culture medium during the hydrogel neutralization step. For all cell culture experiments, hypoxic hydrogels were prepared by swelling 1-part 2% (wt/v %) granular hydrogel with 1-part 2x DMEM culture medium (ThermoFischer Scientific) containing 1 I.U Oxyrase^TM^.

### Microfluidic device design, fabrication and assembly

The microfluidic co-culture device was designed using AutoCAD (version 2020, Autodesk, USA). The device consisted of a central cell culture chamber (6 mm width x 5 mm length x 250 µm height), which was flanked by 2 peripheral culture media chambers (3 mm width x 5 mm length x 250 µm height) separated by a pair of micro-pillar array (300 µm width x 500 µm length and 250 µm height) with 500 µm gaps between the pillars.

The device mold was 3D printed with PlasClear v2.0 (ASIGA, Australia) using ASIGA UV Max X27 printer (ASIGA, Australia) with layer thickness kept at 100 µm. The exposure time for the resin was 13 seconds. The 3D printed mold was first rinsed with isopropyl alcohol (IPA), followed by soaking the mold in IPA for 2 hours to remove excess uncured resin before rinsing with distilled water, air drying and UV curing for 30 minutes. The mold was silanized with trichloro(1H,1H,2H,2H-perfluorooctyl) silane in a vacuum desiccator for 2 hours.

The microfluidic device was then fabricated by replica molding polydimethylsiloxane (PDMS) (Dow Corning, USA) on the 3D printed molds at 10:1 elastomer:curing agent ratio. The microfluidic device was capped with a 250 µm thick PDMS substrate using plasma bonding (at 50 W, 20 sccm of O_2_, for 50 seconds) (FemtoScience, USA), according to a previously established protocol [11]. All seeding inlets and outlets of the device were connected to 1-way stopcocks with Luer connections (Cole-Parmer, USA) with 1.65 mm OD Optimum^®^ 90° angled tips (Nordson EFD, USA). The peripheral media reservoirs were created using a 6 mm OD biopsy punch (Robbins Instruments, USA). Prior to assembly, all parts were sterilized by autoclaving at 121°C for 20 minutes except for the stopcocks, which were first soaked with 70% ethanol for 1 hour before rinsing with sterile 1× PBS and drying before use.

### Computational numerical simulation of oxygen mass transport across granular hydrogel

To simulate the oxygen concentration gradient within the granular hydrogel volume in the microfluidic co-culture device, the lattice Boltzmann method with D2Q5 model was used for the convection-diffusion equation [12]:

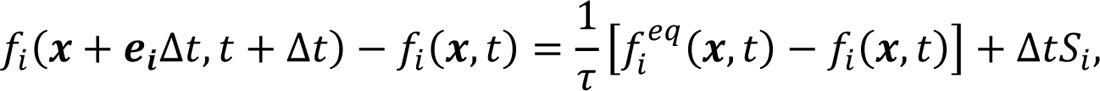

where *f*_*i*_ is the microscopic oxygen concentration distribution function (also called density distribution function) for the *i*-th component at location *x* and time *t*; *e*_*i*_ is the discrete velocities with *e*_0_ = (0,0) and 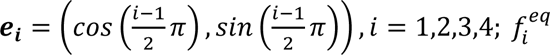 is the equilibrium distribution function, which can be calculated by:

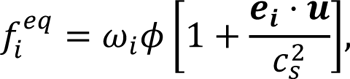

where ω_*i*_ are the weights, being ω_0_ = ⅓ and ω = 1/6 for *i* = 1,2,3,4. The macroscopic scalar ф (oxygen concentration in the current study) can be obtained from 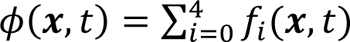. The diffusivity can be controlled by the relaxation parameter, τ by *D* = ⅓(τ − 0.5). More details regarding the method can be found in [12]. (τ − 0.5). More details The simulation was initialized with oxygen concentration of 19.52 g m^-3^ and 0.293 g m^-3^ on the left and right cell culture chambers, respectively. The oxygen diffusivity in Carbomer 940-based granular hydrogel was set to be 4 x 10^10^ m^2^ s^-1^ based on previous literature [13]. Constant concentration boundary conditions were adopted. An oxygen scavenging rate of 1% s^-1^ was used for 1 I.U. Oxyrase^TM^ based on manufacturer’s information within the right chamber to simulate the reaction process.

### Oxygen concentration measurements

Oxygen concentrations in the granular hydrogels, measured as percentage air saturation, were measured using PreSens oxygen profiling microsensor (PM-PSt7-02-L2.5-TS-NB40/0.8-NOP, PreSens GmbH, Germany). Prior to all measurements, the oxygen microsensor was calibrated following the manufacturer’s recommendation. All reading and oxygen concentration measurements were performed under 37°C and 5% CO_2_ similar to the tissue culture operation condition. The microsensor was placed in position for 30s for a stable signal before all measurements were recorded. For on-chip oxygen level measurements, circular access ports of 1.0 mm diameter were created at the centers of each cell culture chambers and the chamber interface using a 1.0 mm biopsy punch (Kai Medica Inc., Japan). Three microsensor access ports were created at 1.5 mm interval from the pillar array, which were located at the center of the normoxic culture chamber, normoxic-hypoxic chamber interface, and center of the hypoxic chamber. Each access port was fitted with separate microsensors. The readout was measured at 24-hour intervals.

### Isolation and culture of primary cells from human cartilage and subchondral bone

Human ethical approval for this project was granted by the Queensland University of Technology (QUT) and the St Vincent Private Hospital Ethics Committees (No. EC00324), and informed consent was obtained from all participating subjects. Briefly, subchondral bone was harvested from patients with primary total knee replacement surgery from tibia surface. Non-sclerotic osteoblasts (nsc-SBO) and sclerotic osteoblasts (sc-SBO) were harvested from the non-sclerotic and sclerotic regions of subchondral bone tissue obtained from a single patient (age 60, male, total knee replacement surgery) following previously established protocol [14]. The subchondral bone samples were treated with collagenase Type II (Thermo Fisher Scientific, Australia) before washing with 1× PBS 3 to 4 times to remove the debris. Isolated osteoblasts were plated in a tissue culture flask and cultured in low glucose DMEM medium supplemented with 10% fetal bovine serum (FBS) and 1% penicillin-streptomycin in a 37°C, 5% CO_2_ incubator. Patient chondrocytes (age 69, male, total knee replacement surgery) were isolated from visually undamaged areas using collagenase Type II (Thermo Fisher Scientific, Australia) as per established protocol [14].

### Chondrocyte spheroid formation and culture

Human primary chondrocyte spheroids were formed by seeding the cells into AggreWell^TM^400 microwell culture plate (STEMCELL Technologies, Canada) at 300, 000 cells per well before incubation at 37^°^C at 5% CO_2_ for 24 hours. To culture the chondrocytes spheroids in granular hydrogels, liquid culture medium was first removed after spheroid formation. The hydrogels were then gently laid over the spheroids in the AggreWell^TM^400 microwell plate before 4-day incubation at 37°C at 5% CO_2_ with 2-day half-hydrogel change intervals. Control experiment of chondrocyte spheroids in liquid DMEM culture media was established. In this configuration, formed spheroids were not harvested from the AggreWell^TM^400. Culture medium was first removed and replaced with fresh culture medium before 4-day incubation at 37°C at 5% CO_2_ with 2-day half-culture medium change intervals.

### Cell viability measurement of chondrocytes in granular hydrogels

To assess cell viability, the liquid DMEM culture medium and the hydrogel overlaying the cultured chondrocytes were first removed before introducing 300 µL of CellTiter 96^®^ Aqueous One Solution Cell Proliferation assay (Promega, Australia) working solution, prepared at 5X dilution in low glucose DMEM culture medium. After 1-hour incubation, the solutions were collected, and read at 490 nm absorbance using a microplate reader (POLARstar, BMGLabtech, Australia). The absorbance values were normalized to 2D cultures containing 300, 000 chondrocytes seeded into 24-well plates maintained under DMEM culture medium for the same duration.

### Patterning and maintenance of cell-laden hydrogel in microfluidic devices

To investigate the effectiveness of laminar patterning of the hydrogels, cell suspension of chondrocytes in hydrogels were used instead of spheroids. 2 paralleled batches of 300, 000 chondrocytes were harvested and pelleted under 20 ×g centrifugation for 1 min. The supernatant was then removed and replaced with 1 mL of serum-free, low glucose DMEM culture medium that was separately treated with CellTracker Blue (Thermo Fisher Scientific, Australia) and CellTracker green (Thermo Fisher Scientific, Australia) for 45 mins under 37°C, 5% CO_2_. After incubating with the CellTracker-treated DMEM culture media, the chondrocytes were pelleted again under 20 ×g centrifugation for 1 min. The supernatant was removed and replaced with 1 mL of normoxic hydrogel. Gentle pipetting of the chondrocytes was performed to resuspend the cells in the hydrogel before loading into a pair of syringes.

The syringes with cell-laden hydrogels were then connected to the stopcocks connecting to the cell seeding inlet of the chips before mounting a dual syringe pump (Legato 111, KDS Scientific). The stopcocks at the cell seeding inlets and outlets were then opened before starting perfusion of the cell-laden hydrogels a flow rate of 0.08 mL/hr until the cell culture chambers were filled. After which, all stopcocks were closed, and the peripheral chambers were filled with their corresponding hydrogels with pipetting. The devices were then immediately image under 364/450 nm excitation/emission wavelength and 488/520 nm excitation/emission wavelength with Nikon Eclipse Ti-2 fluorescence microscope (Nikon, Australia) for cells labelled with CellTracker blue and CellTracker green respectively. Each cell’s position relative to the center within the microfluidic chamber was measured using ImageJ (NIH, USA) via the prebuilt cell tracking plugins.

### Formation of chondrocyte spheroid laden- and osteoblast laden-hydrogel

For cell seeding, the spheroids were harvested post 24-hour incubation and collected in a microcentrifuge tube. The spheroid suspensions were incubated at 37°C, 5% CO_2_ for 10 mins to pellet the spheroids. The supernatants were removed before resuspended in 1 mL of hypoxic hydrogels and normoxic granular hydrogels separately by gentle pipetting with a piston pipette. The spheroid suspended hydrogels were carefully loaded into syringe for seeding. Cell densities were kept at 300, 000 cells per mL. 300, 000 harvested osteoblast were pelleted under 20 ×g centrifugation for 1 min before removing the supernatant. 1 mL of normoxic hydrogels were introduced to resuspend the osteoblast in the hydrogel via pipetting. The osteoblast suspended hydrogels were then loaded into a syringe carefully for seeding.

### Seeding and culture of chondrocyte-spheroids- and osteoblast-laden hydrogels in microfluidic device

A pair of syringes with the chondrocyte spheroid laden hydrogel and osteoblast laden hydrogel were first connected to the stopcocks positioned at the device seeding inlet. The syringes were then mounted onto the dual syringe pump (Legato 111, KDS Scientific). The stopcocks at the cell seeding inlets and outlets were then opened before starting perfusion of the cell-laden hydrogels a flow rate of 0.08 mL/hr until the cell culture chambers were filled. After which, all stopcocks were closed, and the peripheral chambers were filled with their corresponding hydrogels with pipetting. The devices were then placed in a sterile secondary container before incubating in 37°C at 5% CO_2_. Daily replacement of media was performed by pipetting the hydrogels located at the peripheral chambers.

### Immunohistochemistry

Chondrocytes spheroid samples (at approximately 3600 spheroids per mL hydrogel) in the microfluidic devices were harvested by cutting the device and pipetting the cell laden hydrogels into a 1.5 mL microcentrifuge tube. The spheroid samples were washed with 1× PBS 3 times and pelleted via centrifugation using a microcentrifuge. Cell pellets are then fixed with 4% paraformaldehyde (PFA) for 30 mins, permeabilized with 0.5% Triton X-100 in 1× PBS for 30 mins and blocked with blocking buffer (2% BSA/PBS) overnight at 4°C. The samples were then incubated with primary antibodies overnight at 4°C followed by secondary antibodies and DAPI for 2 hours at room temperature. A list of antibodies used is presented in the Supplementary Table 1 and 2. The samples were imaged using Nikon ECLIPSE Ti-E fluorescent microscopy (Nikon, Australia). All images were taken at 300 ms of exposure time to ensure consistency.

### Gene expression quantification

Using a clean blade, the devices were cut in the middle and the cell-laden hydrogels were pipetted into individual 1.5 mL microcentrifuge tube. The cells were then washed with 1× PBS 3 times before pelleting via microcentrifugation to separate the hydrogels from the cells. The cell lysates were then obtained by adding 350 µL of cell lysis buffer (Qiagen, Germany) to the centrifuged cells. Total RNA (tRNA) from the cell lysate were extracted immediately using Qiagen RNeasy Microkit (Qiagen, Germany) before reverse transcription into cDNA using QuantiTect Reverse Transcription Kit (Qiagen, Germany). qPCR was performed with PowerTrack^TM^ SYBR Green Master Mix kit (ThermoFisher, Australia) using Real-Time PCR (Polymerase Chain Reactions) System (ThermoFisher Scientific, Australia). A list of primers (Integrated DNA Technologies, Singapore) for the qPCR is provided in the Supplementary Table 3.

### RNA Sequencing of primary chondrocytes

Total RNA from the nonsclerotic (nsc) and sclerotic (sc) osteoblasts were isolated using Qiagen RNeasy Microkit (Qiagen, Germany) based on the manufacturer’s protocol. Small RNA cDNA libraries were generated with M-MLV Reverse Transcriptase (Promega, M1705) before applying to the Illumina HiSeq2500 (Illumina, USA) 50-nt single-end sequencing. For each sample, the RNA readings were normalized and scaled to counts per million. Statistical analysis and clustering were performed in R (The R Foundation, USA) via the DESeq2 algorithm and p-heatmap functions (|log2Ratio| ≥ 2, FDR ≤ 0.001 were taken as significant). The RNA readings were analyzed using NCBI GenBank (NIH, USA).

### Image quantification

All microscope images were processed as 16-bit images for consistency. Type I and Type II collagen deposition, and HIF-1α activation were quantified by their respective fluorescence intensities normalized by spheroid projected area demarcated by DAPI staining following previous established protocols using ImageJ (NIH, USA).

### Statistical analysis

All results were obtained from at least 3 independent experiments with values expressed as mean ± standard deviation (S.D.). One-way ANOVA was computed for MTS cell viability assays. Two-way ANOVA was computed for normalized HIF-1α intensities across different culture groups with post-hoc Tukey test. Student’s t-tests were computed for gene expressions and normalized collagen intensities for different culture groups. All statistics were calculated using GraphPad PRISM 8.0 (GraphPad, USA) with p<0.05 denoting a significant difference. All graphs were plotted on GraphPad PRISM 8.0 (GraphPad, USA).

## Results

### Generation of hypoxic hydrogels for controlling oxygen concentration

Shear-yielding granular hydrogels can be formed by swelling commercial polyacrylate (Carbomer^®^ 940) microgels in aqueous solution, such as culture media (Fig. 1A, B). The high number of carboxylate groups in the polyacrylate microgels enables hydrogen bonding with water molecules [15], facilitating rapid absorption and swelling in water. These intermittent hydrogen bonds can be disrupted under external forces, resulting in shear-yielding properties of the granular hydrogel [16]. It is important to note that properties of the aqueous solution, including pH, water content, and ionic strength, can affect the water absorption, swelling and stiffness of polyacrylate microgels into a jammed-packed state to form the granular hydrogel (SI Fig. 1) [16, 17]. The polyacrylate microgels swelled maximally in DI water at pH 7 [18] to form a jammed-packed solid without the need for additional crosslinkers (Fig. 1B).

**Figure 1:**
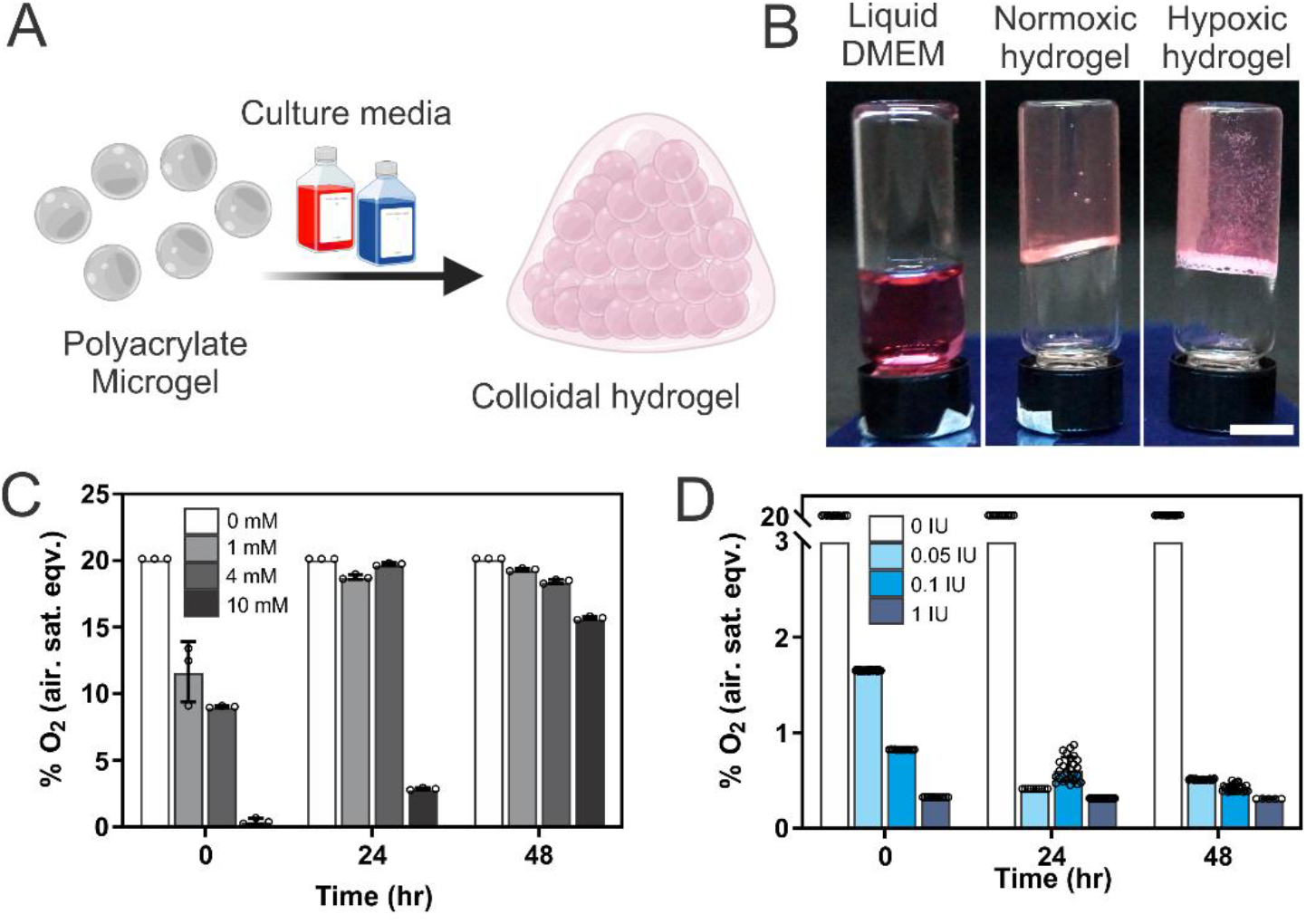
Formulation of hypoxic granular hydrogels. (A) Schematic illustrating the formation of granular hydrogels when polyacrylate microgels were swelled and jammed packed in liquid culture medium. (B) Photos showing successful formation of hypoxic (DMEM + oxygen scavenger) and normoxic (DMEM only) granular hydrogels. Scalebars = 0.6 cm. (C-D) Oxygen concentrations in granular hydrogels containing either (C) sodium sulfite or (D) Oxyrase^TM^ as the oxygen scavengers. Data are average ± SD of technical repeats across 3 separate experiments. Illustration created by Biorender.com

Granular hydrogels formed using culture media have been previously shown to support 3D cell cultures [19] and bioprint soluble growth factors [10]. In this study, we formulated 1% (wt/v) granular hydrogels by mixing equal parts of a 2% granular hydrogel, which was pre-formed and neutralized in DI water, with 2× DMEM culture medium to avoid any pH fluctuation that can affect labile biomolecules in the medium. Using this strategy, both normoxic and hypoxic hydrogels can be formulated by the addition of either chemical (i.e., 4 mM sodium sulfite) or enzymatic (i.e., 1 I.U. Oxyrase^TM^) oxygen scavengers, which are commonly employed in anaerobic bacterial culture [20–23], in the culture medium (Fig. 1B).

We first investigated the oxygen scavenging capacity of different hypoxic hydrogels in a conventional 37°C, 5% CO_2_ cell culture incubator over a 48-hour period (Fig. 1C-D). Control normoxic hydrogels formulated with culture medium without oxygen scavengers recorded an average oxygen concentration of 20.136 ± 0.07% over 48 hours, which corresponded to atmospheric oxygen concentration. Sodium sulfite is a chemical oxygen scavenger that reduces oxygen concentration when sulfite ions irreversibly react with oxygen [24]. As a result, the lowest oxygen level that could be achieved and maintained was directly limited by the concentration of sodium sulfite in the hypoxic hydrogel. Oxygen concentration in the hypoxic hydrogel reverted to ambient levels (19.33 ± 0.08% and 18.46 ± 0.08%, respectively) within 24 hours at lower concentrations of 1- and 4-mM sodium sulfite, likely due to sulfite ion depletion from the chemical reaction (Fig. 1C). Although hypoxic hydrogel with 10 mM sodium sulfite was effective in maintaining oxygen levels at below 5% for 24 hours (Fig. 1C), it lost its oxygen scavenging capacity after 48 hours, resulting in an oxygen concentration of 17.47 ± 0.19% (Fig. 1C). Next, Oxyrase^TM^, an enzyme that catalyzes the reaction of oxygen to form water [20, 23], was incorporated into the hypoxic hydrogel to scavenge oxygen. We observed that hypoxic hydrogels containing Oxyrase^TM^ exhibited concentration-dependent oxygen scavenging capacity within the first 24 hours of incubation. However, this dependency diminished over time, and by 48 hours, an oxygen concentration of ∼0.5% was achievable at all enzyme concentrations (i.e., 0.05, 0.1, and 1 I.U.) tested. Notably, the hypoxic hydrogel with 1 I.U. Oxyrase^TM^ was the most effective in generating and maintaining a hypoxic environment of 0.51 ± 0.01% for 48 hours (Fig. 1D). This period is suitable for tissue culture applications, as it falls within the routine media change interval. Therefore, we employed hypoxic hydrogels with 1 I.U. Oxyrase^TM^ for all subsequent cell culture work.

### Patterning differential oxygen environments with hypoxic hydrogels

Next, we sought to use a microfluidic device to spatially pattern the normoxic and hypoxic hydrogels to control the local oxygen concentration that cells cultured in the device would be subjected to. Microfluidic devices containing multiple chambers separated by micropillars or having different chamber heights enable the patterning of extracellular matrix-based hydrogels, such as collagen and gelatin methacrylate (GelMA), via capillary and surface tension forces [25, 26]. Similarly, we employed a microfluidic device that consisted of a central cell culture chamber, which was flanked by two reservoir chambers separated by micropillar arrays (Fig. 2A). By leveraging on the shear yielding and viscous properties of the granular hydrogels, we could easily achieve laminar flow patterning of two different cell-laden hydrogel streams in the central cell culture chamber (Fig. 2Ai). The cells were fed by routine replenishment of fresh granular hydrogels swelled in culture media loaded into the periphery reservoir chambers. The micropillar arrays between the cell culture and reservoir chambers acted as barriers to minimize disturbances to the patterned cell-laden hydrogels during daily hydrogel replenishment (Fig. 2Aii). Using this patterning strategy, we showed that two populations of chondrocytes (pre-labelled with Cell Tracker Green and Blue) embedded in the granular hydrogel can be as effectively patterned (Fig. 2Bi, C).

**Figure 2:**
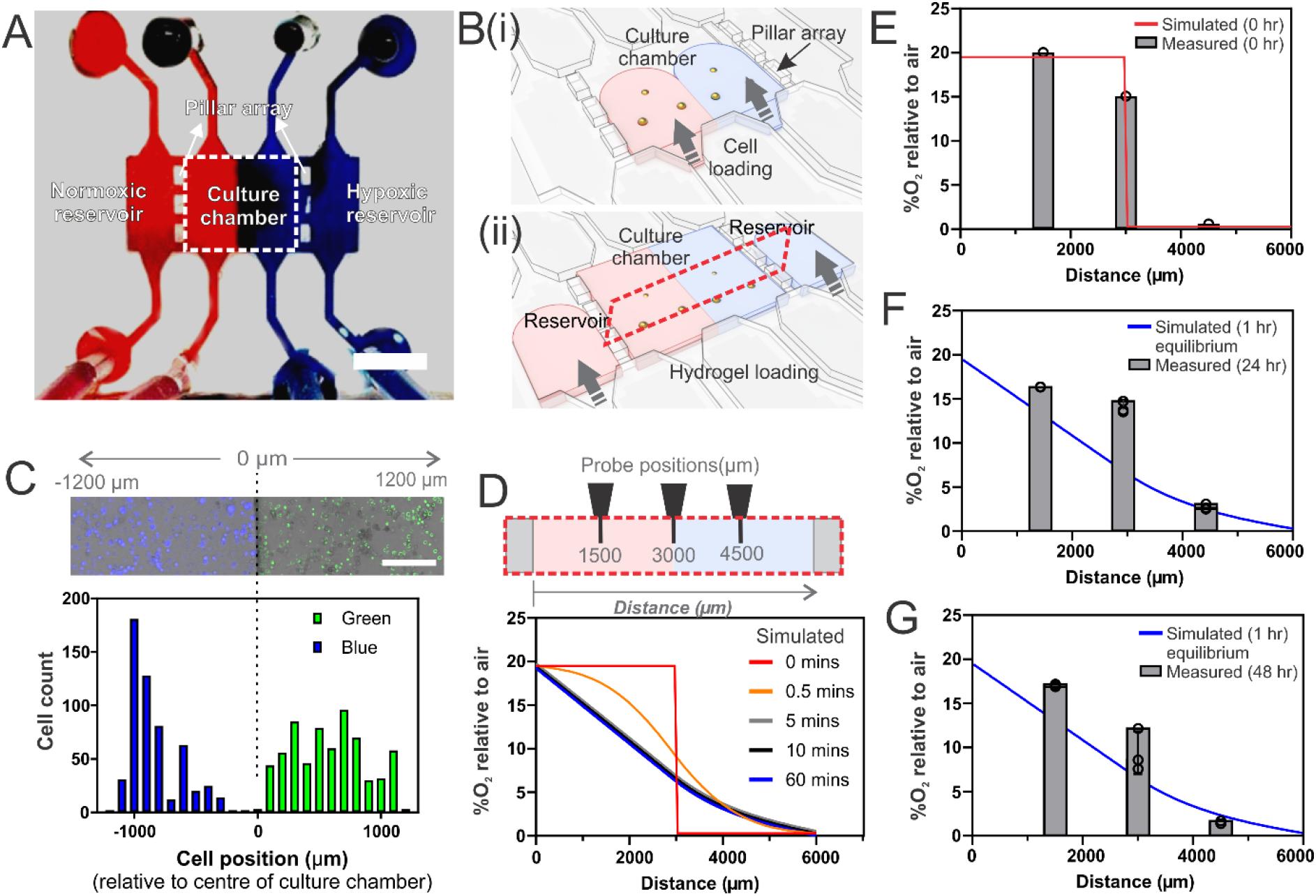
Spatial patterning of cells and oxygen concentration within a microfluidic device. (A) The microfluidic device features a central cell culture compartment, flanked by two reservoir compartments, and separated by micro-pillar arrays. This design allows for (i) the laminar flow-based seeding of cells in dual streams of normoxic (indicated in red) and hypoxic (indicated in blue) hydrogels and (ii) nutrient replenishment through hydrogel replacement in the reservoir chambers, all with minimal disturbance. (B) Cross-sectional schematics of the cell culture compartment illustrate (i) cells patterned and fluorescently labeled in blue and green (scale bar = 150 μm), and (ii) the locations of oxygen-sensing probes. (C) A histogram quantifies the distribution of blue and green-labeled cells across the width of the cell culture compartment, noting minimal cell mixing at the chamber interface (designated as 0 µm within the device). (D) Computational simulations reveal the time-dependent establishment of oxygen gradients post-patterning with normoxic and hypoxic hydrogels. (E-G) Experimental measurements of oxygen concentration at three distinct locations in the cell culture compartment (as detailed in B, ii) are shown at different post-incubation timepoints at 37°C, 5% CO2: (E) 0 hours, (F) 24 hours, and (G) 48 hours. Data represents the mean ± S.D. from three independent experiments.

To estimate the formation of a steady-state oxygen gradient across the hydrogel, we conducted analytical modeling using the diffusion coefficient of oxygen in Carbomers solution obtained from literature [13]. Due to the rapid enzymatic activity of Oxyrase within the small hydrogel volume, a steady-state oxygen gradient was established within 60 minutes of incubation at 37°C (Fig. 2D). This rapid establishment of an oxygen gradient is advantageous as it minimizes changes in the culture microenvironment. We then experimentally characterized the oxygen concentration gradient that resulted from patterning normoxic and hypoxic hydrogels formulated with 1 I.U. Oxyrase^TM^ in the microfluidic device. The oxygen levels at the center of the normoxic region, hypoxic region and their interface were measured with an oxygen microsensor probe immediately after hydrogel patterning (0 hour) and after 48 hours of incubation in a 37°C, 5% CO_2_ cell culture incubator (Fig. 2E-G). We observed sustained oxygen gradient across the microfluidic cell culture chambers, which concurred well with the simulated equilibrium concentration gradient (Fig. 2D).

### Chondrocyte cellular responses in hypoxic hydrogels

We then investigated whether the granular hydrogel is biocompatible and supported physiological cell phenotypes and functions. Primary human chondrocyte spheroids were employed as the putative cell model since it is well established that chondrocytes exhibited more physiological-relevant phenotypes when cultured in a hypoxic environment with <5% oxygen [27–29]. We first measured cell viability of 3D chondrocyte spheroids cultured in hypoxic and normoxic hydrogels, with and without 1 I.U. Oxyrase^TM^ respectively, using the MTS assay after 4 days of culture. It was observed that there was no significant difference in the viability of the 3D chondrocyte spheroids in both normoxic and hypoxic hydrogels as compared to the 2D control (Fig. 3A). This observation concurred with existing literature on the biocompatibility of polyacrylate-based granular hydrogels [9, 19, 30].

**Figure 3:**
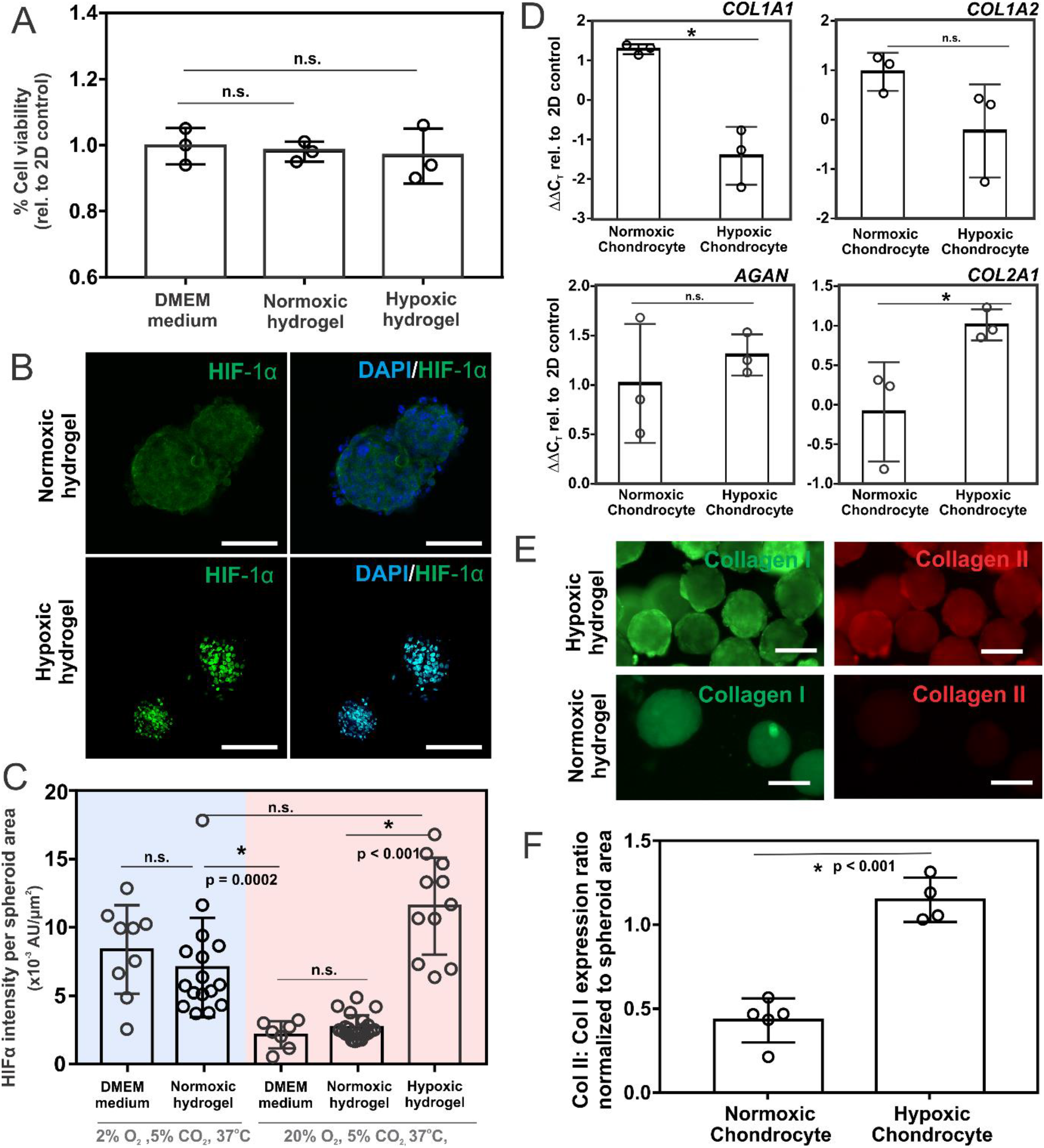
Viability and hypoxia-responsive behavior of primary human chondrocyts in granular hydrogels. (A) Viability assays for primary human chondrocyte spheroids cultured in liquid medium, normoxic, and hypoxic hydrogels over a 4-day period. Each data point corresponds to individual, separate, experiment. (B) Immunofluorescent visualization of HIF-1α expression in chondrocyte spheroids following 4 days in normoxic and hypoxic hydrogel environments (scale bar = 50 μm). (C) Quantitative analysis of HIF-1α expression levels in chondrocytes cultured in liquid DMEM medium, normoxic, and hypoxic hydrogels under either conventional (20% O_2_, 5% CO_2_) or hypoxic (2% O_2_, 5% CO_2_) incubator conditions. HIF-1α fluorescence intensities are normalized to the projected spheroid area delineated by DAPI nuclear staining. (D) Gene expression profiles of *COL1A1, COL1A2, AGAN*, and *COL2A1* after 4 days in normoxic and hypoxic hydrogels. (E) Immunofluorescent imaging of Type I (green) and Type II (red) collagen following 4 days in normoxic and hypoxic hydrogels (scale bar = 100 μm). (F) Quantitative comparison of relative intensities for Type II to Type I collagen expression in chondrocyte spheroids, normalized to spheroid area as identified by DAPI staining. Asterisks (*) indicate statistical significance: p<0.05, evaluated via (C) one-way ANOVA with post-hoc analysis and (F) Student’s t-test.

We then assessed whether the cellular adaptation response in the hypoxic hydrogel was similar to cells cultured in a conventional hypoxic incubator maintained at 2% O_2,_ 5% CO_2,_ 37°C. Cell adaptation to hypoxia is directly regulated by the oxygen sensitive master transcription factor, hypoxic inducing factor 1 (HIF-1α). Hence, we assessed HIF-1α expressions in the primary chondrocyte spheroids when they were cultured in liquid medium or granular hydrogels placed in incubators with oxygen levels set at either 20% or 2% (Fig. 3B, C). In a hypoxic incubator, it was observed that HIF-1α had translocated into the cell nuclei regardless of whether the chondrocytes were maintained in liquid culture medium or granular hydrogels (Fig. 3B, SI Fig 2), indicating that cells grown in the granular hydrogel could respond to pre-set oxygen level in the incubator. Inside a conventional 20% O_2,_ incubator, increased expression and nuclear translocation of HIF-1α were only observed when the chondrocytes were maintained in hypoxic granular hydrogels (Fig. 3B). In comparison, HIF-1α expression was dispersed (Fig. 3B) and at significantly lower levels (Fig 3C) (one-way ANOVA, p=0.0002) when chondrocytes were cultured in either liquid culture medium or normoxic hydrogels without Oxyrase™ (Fig. 3C). This result is consistent with the low oxygen concentration measured in the hypoxic hydrogel (Fig. 1D) and demonstrates that we could elicit a hypoxic cellular response without the need for hypoxic incubators.

We then aim to investigate the influence of oxygen concentration on cartilage-specific phenotypes. Given that cartilage tissue is usually avascular, chondrocytes are naturally exposed to hypoxic conditions [27]. Previous research has demonstrated that hypoxic culture conditions increase Type II Collagen expression, resulting in a more anabolic phenotype that is associated with hyaline cartilage. On the other hand, normoxic culture conditions favour the production of Type I Collagen, leading to a more catabolic phenotype associated with fibrous cartilage [31, 32].

To examine the primary chondrocyte’s response to changes in oxygen levels, we performed 3D culture of chondrocyte spheroids in hypoxic and normoxic hydrogels, comparing them to standard 2D cultures of chondrocytes in normoxic 24-well plates. We assessed the expression levels of genes known to affect fibrous (*COL1A1, COL1A2*) and hyaline (*AGAN, COL2A1*) cartilage phenotypes using RT-qPCR (Fig. 3D), and our data supported previous findings based on tissue histology [31, 32]. Specifically, we observed a significantly lower expression level of *COL1A1* in hypoxic hydrogel cultures while expression level of *COL2A1* was significantly increased (Fig 3D). The higher expression of *COL2A1* under hypoxic hydrogels may be attributed to the role of HIF-1α in type 2 collagen synthesis [33]. Next, we compared the relative protein expression of Type I and Type II collagen measured by immunohistochemistry in chondrocyte spheroids cultured using hypoxic and normoxic hydrogels (Fig 3E). For chondrocyte spheroids cultured in hypoxic hydrogels, we noted a significantly higher ratio of Type II to Type I collagen (Fig. 4F), which agreed with our gene expression results. In all, our data indicated that the hypoxic granular hydrogel could maintain primary chondrocytes to exhibit a more physiological phenotype typically observed in low-oxygen environment without using hypoxic incubators.

**Figure 4:**
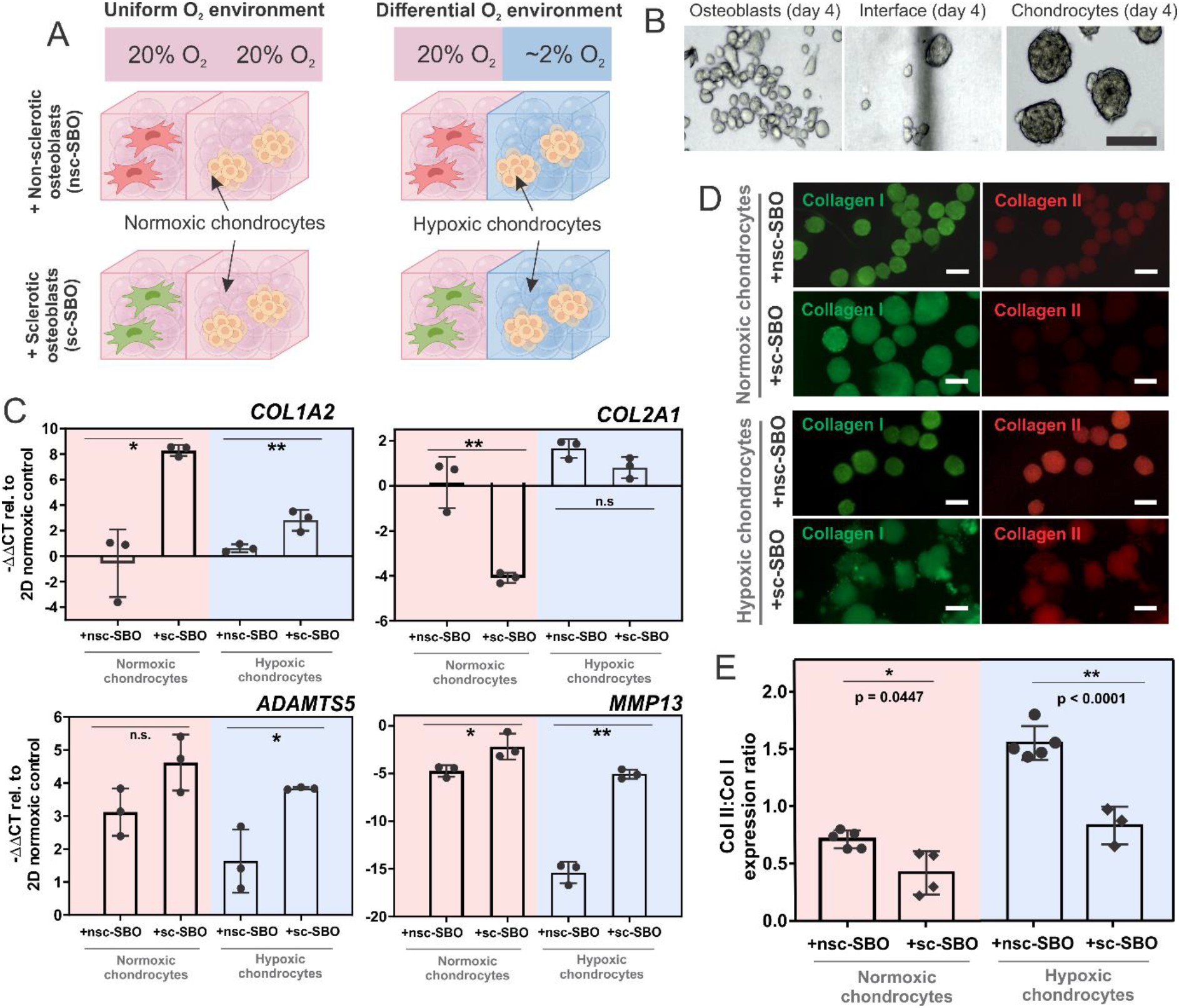
Microfluidic co-culture of chondrocytes and osteoblasts under differential or uniform oxygen tensions. (A) Experimental design illustrating chondrocytes paired with both non-sclerotic (nsc-SBO) and sclerotic (sc-SBOs) osteoblasts under different oxygen conditions. (B) Day 4 light microscopy images showing co-cultured osteoblasts and chondrocyte spheroids embedded in normoxic and hypoxic hydrogels, respectively. (Scalebar = 150 μm). (C) RT-qPCR analysis of *COL1A2, COL2A1, ADAMTS5*, and *MMP13* gene expression in both uniform and differential O_2_ co-cultures. (D) Immunohistochemical staining depicting Type I (green) and Type II (red) collagen in chondrocytes in both uniform and differential O_2_ co-cultures with nsc-SBOs and sc-SBOs. (Scalebar = 100 μm). (E) Intensity quantification of Type II:Type I collagen in chondrocytes. Asterisks denote statistical significance: *p<0.05, **p<0.001, via Student’s t-test. Schematic in (A) created with Biorender.com.

### Osteoblast-chondrocyte coculture under differential oxygen tension

It is well established that OA progression is characterized by cellular changes in both cartilage and the underlying subchondral bone, and crosstalk mechanisms exist between cells in these tissues [34–36]. Studies by our team and others have indicated that one such crosstalk mechanism involves the induction of catabolic effects in chondrocytes by proinflammatory soluble factors produced by osteoblasts from sclerotic subchondral bone as a result of increased mechanical loading during OA [14, 37, 38]. Hence, there have been active research efforts to develop therapeutic strategies targeting the normalization of one cell type, which could reverse or slow down OA progression in both bone and cartilage [39, 40]. Hence, in vitro OA disease wmodels designed to functionally assess potential therapeutic strategies would need to recapitulate the osteochondral interface tissue as a physiological functional unit. Since we showed that the hypoxic granular hydrogel could maintain primary human chondrocytes to exhibit a more anabolic phenotype independently of hypoxic incubators, we attempted to generate a microfluidic osteoblast-chondrocyte co-culture model that reflects the differential oxygen environment that exists in the osteochondral interface tissue by patterning normoxic and hypoxic cell-laden hydrogels in a microfluidic device.

Allogenic pairs of primary human osteoblasts and chondrocytes were used in the microfluidic co-culture model to study osteoblast effects on chondrocyte phenotype (Fig. 4A). A control co-culture in normoxic hydrogels was also established. Sclerotic (nsc-SBO) and non-sclerotic osteoblasts (sc-SBO) were sourced from patient subchondral bone zones [14]. Previous RNA sequencing confirmed significant differences in exosome profiles between nsc-SBO and sc-SBO, notably affecting Type II collagen expression in chondrocytes [14]. In the bone-cartilage co-cultures, primary chondrocytes were in hypoxic hydrogel with 1 I.U. Oxyrase™, and sc-SBO or nsc-SBO in normoxic hydrogels. These were patterned in a microfluidic device as previously shown (Fig. 3A) and cultured in a conventional 20% O_2_, 5% CO_2_ incubator for 4 days. Nutrients were refreshed every 2 days. Both cell types showed good viability in these differential oxygen conditions over the 4-day period (Fig. 4B).

We aimed to assess the influence of healthy nsc-SBO and pathogenic sc-SBO osteoblasts on chondrocyte collagen synthesis and the expression of catabolic enzymes under a differential oxygen level (5% −20%) or a uniform oxygen level (20%). Our findings revealed that coculturing chondrocytes with sc-SBOs significantly increased the expression of Type I collagen (*COL1A2*) by 14.97-fold and 4.87-fold in the differential and uniform oxygen co-cultures, respectively (Fig. 4C). Conversely, sc-SBOs attenuated the expression of Type II collagen (*COL2A1*), although the reduction was more prominent in the uniform oxygen co-culture (29.29-fold) compared to the differential oxygen co-culture (2.06-fold) (Fig. 4C). Interestingly, the impact of sc-SBOs on chondrocytes was more pronounced in inducing catabolic enzymes under the differential oxygen environment (Fig. 4C). Specifically, *ADAMTS5*, an aggrecanase, was upregulated by 1.48-fold (p = 0.133) in the uniform oxygen co-culture (Fig. 4C), while co-culture under differential oxygen levels showed a significant 2.35-fold upregulation (p = 0.026) (Fig. 4C). A similar trend was observed for the collagenase, *MMP13*, where the uniform oxygen co-culture exhibited a 2.14-fold upregulation (p = 0.046), and the co-culture with differential oxygen levels showed a 3.01-fold upregulation (p < 0.0001).

Similarly, we could detect sc-SBO induced cartilage matrix catabolism at the protein level. The relative amount of Type II to Type I collagen (Col II / Col I ratio) in the chondrocyte spheroids were determined when they were co-cultured with either nsc-SBOs or sc-SBOs under uniform or differential oxygen levels (Fig. 4D-E). In co-culture with sc-SBOs, we observed a significant reduction in Col II / Col I ratio, approximately 2-fold lower than nsc-SBO co-cultures, regardless of oxygen conditions (Fig. 4E). This change was more pronounced in the context of differential oxygen levels, where the Type II to Type I collagen ratio was inherently higher in chondrocytes, aligning with earlier findings that hypoxic conditions promote Type II collagen production. We also evaluated the impact of differential oxygen levels on both nsc-SBOs and sc-SBOs in the co-culture by assessing the expressions of osteoblast differentiation markers, alkaline phosphatase (*ALP)* and osteopontin (*SSP1*). No significant differences in *ALP* and *SSP1* expressions were observed in nsc-SBOs, and *ALP* expression in sc-SBOs, regardless of the oxygen conditions (SI Fig. 3). A minor increase in *SSP1* expression was observed in sc-SBOs in hypoxic conditions. The results suggest that oxygen levels had a limited effect on osteoblast phenotype.

Taken together, our results highlighted the effects of sc-SBOs on chondrocyte collagen synthesis and the expression of catabolic enzymes were modulated by oxygen conditions. This suggests a potential role of oxygen levels in modulating osteoblast-chondrocyte crosstalk to induce a catabolic phenotype in chondrocytes.

### Differential oxygen level mimics physiological changes in chondrocyte metabolism induced by sclerotic subchondral osteoblasts

To gain further insights into the mechanistic relationship between oxygen tension and SBO-chondrocyte crosstalk, we performed RNA sequencing to examine the differences in chondrocyte transcriptomes under differential or uniform oxygen conditions during co-culture with nsc-SBOs and sc-SBOs. We first examined how oxygen level affected chondrocyte transcriptome during co-culture with nsc-SBOs. A total of 9423 genes were differentially expressed (DEGs) between the two oxygen conditions (p < 0.05, false discovery rate, FDR < 5%) as visually represented in a Volcano Plot, with many genes involved in the cartilage matrix development (Fig. 5A). Gene ontology analysis revealed a significant enrichment of pathways involved in the ECM matrix organization and collagen-mediated ECM interactions in chondrocytes when they are maintained in hypoxia in a co-culture with differential oxygen level (SI Fig. 4). To confirm if these oxygen-induced DEGs impacted chondrocyte differentiation and metabolism, we first queried the NCBI GenBank database to identify genes associated with cartilage development and cartilage metabolism, and curated a list of genes with well reported regulatory functions in chondrocyte metabolism and are often dysregulated in OA (SI Table 4) [41–44]. It was observed that when chondrocytes were in a normoxic co-culture condition, there was upregulation of cartilage catabolism associated markers. They include hypertrophic differentiation genes (e.g. *VEGFA4, VEGFA9, MMP13 and MMP1)* [45], proinflammatory factors (e.g. *IL6, IL1B*, and *IL17RB)* (Fig. 5B). Conversely, there was downregulation of genes involved in anabolic signaling pathways, such as I*GF1, TGF-β* and *FGF2*, as well as downregulation of aggrecan (*ACAN.3*) (Fig. 5B). These data buttressed our earlier observations in Fig. 3 that chondrocytes maintained in the hypoxic hydrogels better maintained an anabolic phenotype compared to those grown in normoxic hydrogels.

**Figure 5:**
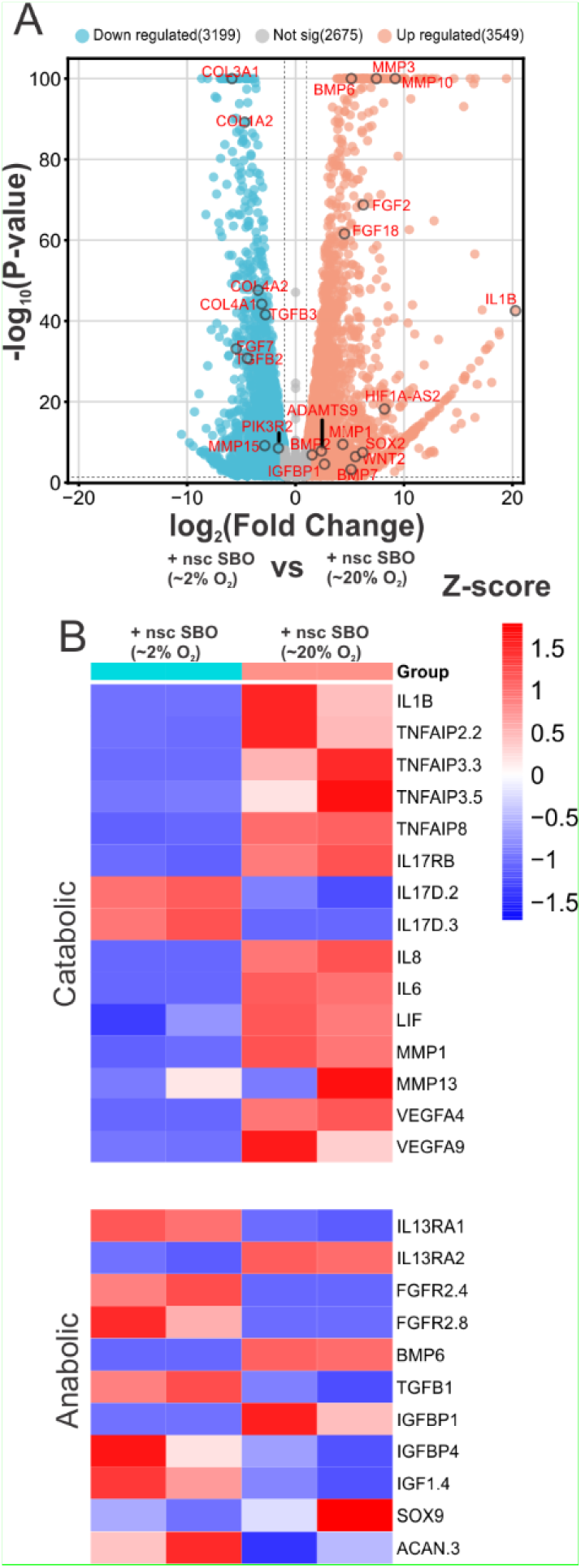
Differentially expressed genes (DEGs) of hypoxic (2% O2) and normoxic (20% O2) chondrocytes co-cultured with nsc-SBOs in the microfluidic device. (A) Volcano plot of the DEGs between hypoxic and normoxic chondrocytes co-cultured with nsc-SBOs. Annotated genes (in red) are associated to chondrocyte cartilage development. (B) Heatmap of DEGs involved in cartilage metabolism for hypoxic and normoxic chondrocytes co-cultured with nsc-SBO. Each column represents individual measurements of n = 3 pooled samples. Only differentially expressed genes with FDR<0.05 were identified.

Next, we conducted a comparative analysis between nsc-SBO and sc-SBO co-cultures under both differential and uniform oxygen conditions. This was to investigate how the metabolic state of chondrocytes may affect the changes induced by pathogenic sc-SBOs. In uniform normoxic co-cultures, fewer DEGs emerged. As compared to nsc-SBOs, sc-SBOs upregulated 681 genes and downregulated 244 genes within 5% FDR (Fig. 6A). Among these, we observed that sc-SBOs induced upregulation of some proinflammatory genes, such as *IL17D* and *IL8*, although most catabolic genes within our curated list remained unchanged. We also observed that sc-SBOs downregulated Insulin Growth Factor-1 (*IGF1*) and Transforming Growth Factor-β1 (*TGFβ1)*, which are the primary anabolic growth factors mediating cartilage differentiation and anabolism (Fig. 6B) [46, 47].

**Figure 6:**
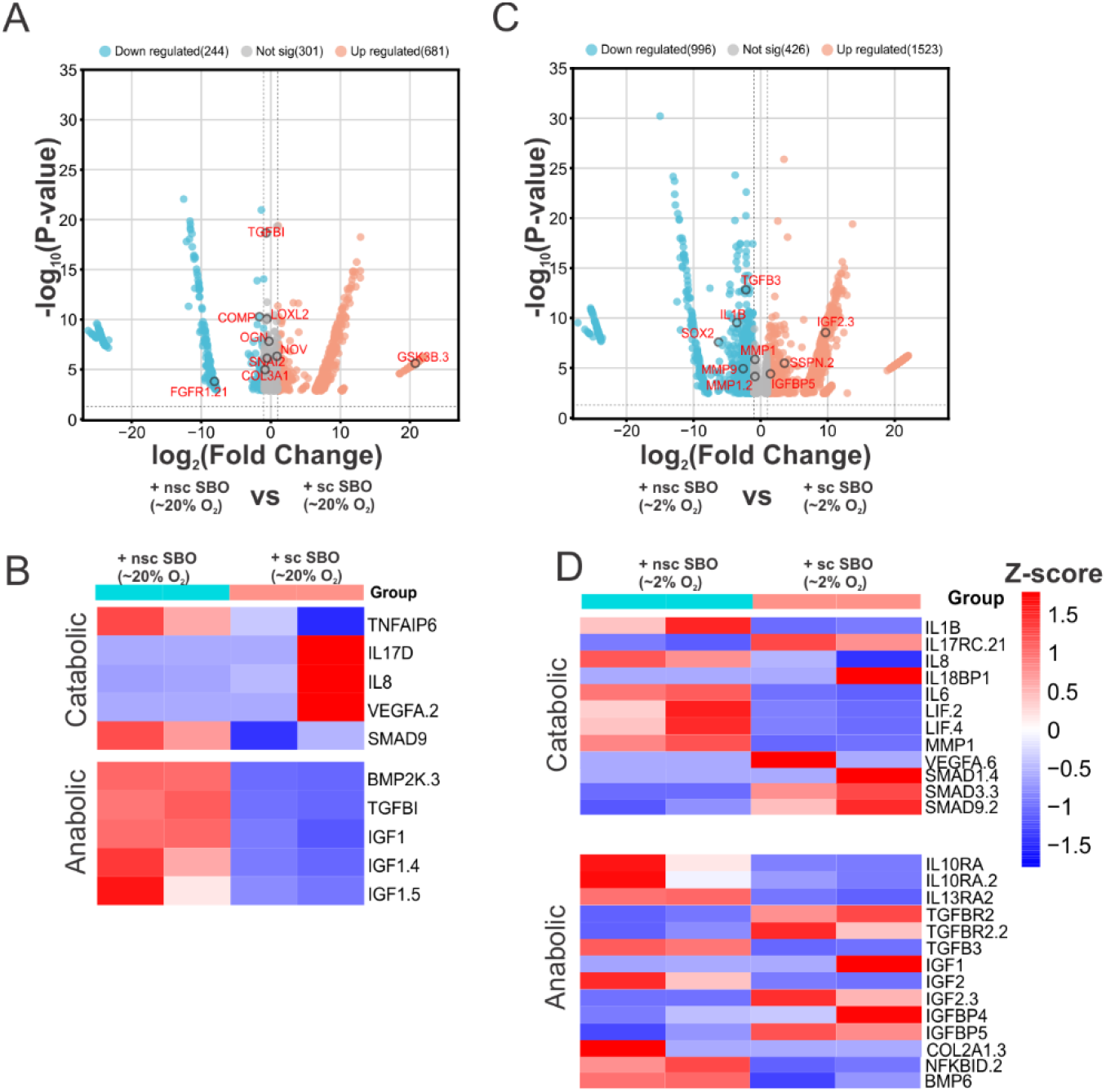
Differentially expressed genes of co-cultured chondrocytes in the microfluidic device between the nsc-SBO and SC-SBOs under normoxia and hypoxia. (A) Volcano plot of the DEGs normoxic chondrocytes co-cultures with nsc-SBOs and sc-SBOs with the labelled genes related to cartilage development in normoxia (B) Heatmap of DEGs in cartilage metabolism for normoxia chondrocyte co-cultures with nsc-SBO. Each column represents individual measurements of n = 3 pooled samples. (C) Volcano plot of the DEGs normoxic chondrocytes co-cultures with nsc-SBOs and sc-SBOs with the labelled genes related to cartilage development in hypoxia. (D) Heatmap of DEGs in cartilage metabolism for hypoxia chondrocyte co-cultures with nsc-SBO. Each column represents individual measurements of n = 3 pooled samples. Only differentially expressed genes with FDR<0.05 were identified.

When we investigated the impact of sc-SBOs on the metabolism of chondrocytes, which were maintained under hypoxia in a co-culture with differential oxygen levels, we observed that the number of DEGs was substantially higher than in a normoxic co-culture, with 1523 upregulated genes and 996 downregulated genes (Fig. 6C). There are also significantly higher number of DEGs associated to cartilage metabolism related to OA observed (Fig. 6D). sc-SBOs led to an increase in transcription of proinflammatory factors, *IL17* and *IL18*, and depressed anti-inflammatory factors, *IL10* and *IL13* (Fig. 6D). All these cytokines are known to directly modulate metalloprotease activity [48]. Interestingly, canonical inflammatory factors of the cartilage, such as *IL1B, IL8* and *IL6* were downregulated. We also noted differential activities in the NF-κβ pathway (Fig. 6D). The downregulation of NF-κβ inhibitor delta (*NFKBID*) together with elevated *IL17* expression suggested activation of canonical NF-κβ signaling, which promotes metalloprotease and matrix catabolism [44]. In hypoxic chondrocytes, sc-SBOs not only influenced IGF and TGF-β/BMP signaling pathways by altering ligand expression observed in the normoxic chondrocytes, but also affected the mediators involved in these networks. For instance, we observed a downregulation of *BMP6* and *TGFΒ3*, which was accompanied by upregulation of its receptor *TGFBR2* (Fig. 6D), suggesting a compensatory mechanism due to lack of ligand availability. The resultant elevated SMAD1/3/9 expressions suggest sc-SBO altered TGF-β/BMP signalings in hypoxic chondrocytes, which may promote terminal differentiation of chondrocytes towards a hypertrophic state often seen during endochrondral ossification and OA [49]. We also observed pronounced changes in the transcription of IGF ligands and IGF binding proteins (IGFBPs), which regulate the bioavailability of IGFs. IGFBPs are often found at elevated levels in OA cartilage, which attenuate the responsiveness of chondrocytes to the anabolic action of IGF ligands [47, 50]. *IGF1* expression was upregulated but not *IGF2* (Fig. 6D). *IGFBP4* and *IGFBP5* transcription was also increased. These results concurred with previous studies profiling these factors in primary cartilage tissues and chondrocytes obtained from OA patients [50, 51]. Taken together, our unique microfluidic co-culture model featuring patterned hypoxic and normoxic hydrogels effectively mimics sc-SBO-induced changes in chondrocyte catabolism and anabolism, an outcome not seen in uniform normoxic co-cultures.

## Discussion

Polyacrylate granular hydrogels offer unique advantages in tissue culture over hydrogels that require chemical, heat, or light crosslinking. These hydrogels form soft solids through intermittent hydrogen bonding between polyacrylate microgels [16]. At pH 7, they achieve maximum swelling and soft solid formation due to a high number of inter-polyacrylate hydrogen bonds [18]. This unique feature allows us to create cell-laden hydrogels with tunable biochemical properties by altering the composition of the aqueous culture medium in which the microgels are swelled. We leveraged this property to generate both normoxic and hypoxic hydrogels by adding oxygen scavengers to the swelling culture medium. Using oxygen scavengers at varying concentrations, we successfully modulated the oxygen levels in the hydrogel (Fig. 1C–D). We also achieved stable, 48-hour oxygen gradients by patterning normoxic (20% O_2_) and hypoxic (<1% O_2_) hydrogel streams side by side in a microfluidic device (Fig. 2). This eliminates the need for complex microfluidic systems traditionally used for gradient control [7, 8]. Unlike the patterning of soluble factors like growth factors, maintaining a dissolved oxygen gradient is more challenging due to ambient oxygen influx. To effectively retain the patterned oxygen levels, the choice of oxygen scavenger is crucial. Different scavengers have varying capacities to deplete oxygen, and their efficacy is directly related to the resulting hydrogel oxygen levels. The duration of this effect is influenced more by the scavenger’s reaction kinetics than by its inherent strength. Our findings indicate that irreversible reducing agents, such as sodium sulfite, are less effective for long-term applications compared to reversible or catalytic agents like Oxyrase™ (Fig. 1C–D).

In addition to chemical composition tunability [19], the granular hydrogels offer adjustable mechanical properties to meet tissue culture demands. By modulating factors, such as hydrogel content [52], pH [16, 53], and ionic content [54], we showed that the hydrogel’s stiffness can be tuned (SI Fig. 3). This versatility potentially allows for fine-tuning the physical properties of the hydrogel to mimic the native tissue microenvironment and optimize responses of multiple cell types more accurately. In the current study, a 1-2% polyacrylate-based granular hydrogel yielded stiffness ranging from 37.8 to 81.6 kPa (SI Fig. 3). This is notably lower than the stiffness found in patient-derived articular cartilage, which is typically in the MPa range [55, 56]. While an increase in polyacrylate microgel content could enhance the hydrogel’s stiffness, additional crosslinking strategies can be employed. For instance, combining polyacrylate with other materials like Poly(N-isopropylacrylamide) (PNIPAm) can significantly elevate the hydrogel’s stiffness to levels comparable with human cartilage [57, 58].

In this study, we have shown that hypoxic granular hydrogels formulated with Oxyrase™ effectively induced hypoxic responses in primary human chondrocytes. This was evident in both monoculture (Fig. 3) and coculture with osteoblasts (Fig. 4-5). Notably, hypoxia-inducible factor 1-alpha (HIF1α), the key regulator of hypoxic responses, displayed increased expression levels and nuclear translocation in chondrocytes cultured in hypoxic hydrogels within a conventional incubator (Fig. 3B-C, Fig. 5A). Elevated nuclear activity of HIF1α was associated with a higher Type II to Type I collagen ratio in chondrocyte spheroids (Fig. 3C). This can be attributed to multiple mechanisms: HIF1α directly activates SOX9 to stimulate chondrocyte differentiation markers such as Type II collagen and aggrecan synthesis [59, 60]. Furthermore, HIF1α has been documented to inhibit Wnt signaling, thereby mitigating catabolic activities, including the production of matrix metalloproteinase 13 (MMP13) [61]. This suppression was confirmed in our hypoxic chondrocytes compared to their normoxic counterparts (Fig. 4C-E, Fig. 5B,). In contrast, chondrocytes in normoxic hydrogels exhibited a catabolic phenotype (Fig. 5B), likely driven by elevated ROS levels that induce catabolic genes like VEGF [62] and MMPs [43]. Importantly, the hypoxic and normoxic granular hydrogels can be easily patterned within a microfluidic device to generate a localized hypoxic microenvironment in a coculture system (Fig. 2), which can be used to probe interactions between physiologically hypoxic chondrocytes and other cell types.

The gradual degeneration of cartilage matrix during OA results from the dysregulation of multiple signaling pathways in chondrocytes, which are involved in the regulation of metalloprotease-mediated matrix degradation and collagen production [44]. Recent evidence suggests the sclerotic state of osteoblasts can impact chondrocyte metabolism and matrix remodeling [34]. Thus, we investigated whether a microfluidic osteoblast-chondrocyte coculture model maintained under differential oxygen level achievable using spatially patterned hypoxic and normoxic granular hydrogels can better recapitulate these osteo-chondral crosstalk mechanisms. We observed that the presence of sc-SBOs altered chondrocyte collagen synthesis (Fig. 4D-E) and metalloproteases expressions (Fig. 4C) at the transcriptional and protein level, which were mediated by transcriptional changes in various upstream regulators for cartilage metabolism (Fig. 6). Compared to normoxic co-cultures, we noted that hypoxic co-cultures captured more observable upstream regulatory pathways involved in chondrocyte’s matrix catabolism during OA, including IL-6, IL-1β, NF-κβ and BMP signaling pathways (Fig. 6). Increased activation of the NF-κβ pathway via elevated *IL-17* and reduction of NF-κβ inhibitors (*NKFBID*) alongside elevated SMAD expression in the TGFβ/ΒΜP signaling pathway can lead to the upregulation of metalloprotease, MMP13, to promote matrix degradation [44, 63].

Besides recapitulating sc-SBO induced changes in chondrocyte matrix catabolism regulatory pathways, the differential oxygen co-culture model can also capture early cartilage damage repair mechanisms during early onset of OA. Among the sc-SBO paired chondrocytes, we noticed a concomitant decrease in *IL-6* and *IL-1β* expressions and increase in *IGFBP* and *IGF-1* expressions (Fig. 6D). It is well-established that IGFs serve as the primary anabolic growth factors in cartilage, playing a crucial role in structural maintenance by stimulating the production of cartilage matrix molecules, including type II collagen [46, 47], through the phosphoinositide 3-kinase (PI3K) pathway [64]. Elevated levels of IGFBPs during OA is known to bind to IGF-1 which inhibited the PI3K pathway. Responding to the lower availability of IGF-1 in early OA, chondrocytes homeostasis increases IGF-1 production to ameliorate cartilage degradation [65] by suppressing proinflammatory IL-6 and IL-1β pathways. Collectively, the differential oxygen co-culture model can recapitulate complex molecular interplay involving both collagen synthesis and degradation. This could have explained why sc-SBOs induced a more significant decrease in Collagen II at the protein level (Fig. 4D-E) even though the extent of decrease in *COL2A* gene expression was much smaller than in the uniform normoxic co-culture model (Fig. 4C). In this study, our 4-day co-culture model can only capture multiple catabolism pathways of cartilage including the responses of chondrocytes in early stages of osteoarthritis. An extended period (more than 21 days) of co-culture under the patterned differential oxygen microenvironment could help establish a more definitive anabolic profile of chondrocytes [66].

## Conclusion

Granular hydrogel formed by swelling polyacrylate microgels in oxygen scavengers can be used as a functional material to exert spatio-temporal control over the oxygen microenvironment of a cell culture without the need for specialized instrumentation. We demonstrate that this functional material can be incorporated into a microfluidic device to pattern human primary osteoblasts and chondrocytes with a physiological differential oxygen gradient that occurs at the osteo-chondral interface. This enabled the co-culture model to recapitulate a greater extent of molecular interplay involved in chondrocyte metabolism, matrix synthesis and degradation as compared to a uniform normoxic co-culture model. The micropatterned hypoxic granular hydrogel can be widely adopted to mimic other tissue interfaces with oxygen gradients to better mimic normal and patho-physiology.

## Acknowledgments

This work is supported Australian Research Council (FT180100157 and DP200101658) awarded to YCT, QUT DVC ECR (Early Career Researchers) grant (323100-0235) awarded to LJYO. This work was enabled by use of the Central Analytical Research Facility (CARF) at the Queensland University of Technology (QUT).

## Author Contributions

Conceptualization and experiment designs were performed by LJYO and YCT. LJYO, ZW and JL conducted experimentations and data collection. Data analysis was performed by LJYO, IP, ARS and YCT. Manuscript writing was performed by LJYO and edited by YCT.

## Notes

### Competing Interest Statement

The authors have declared no competing interest.

