## Supplementary Information for "Localized oxygen control in a microfluidic osteochondral interface model recapitulates bone-cartilage crosstalk during osteoarthritis"

Louis Jun Ye Ong[1-3], Antonia Rujia Sun[1-2], Zhongzheng Wang[1], Jayden Lee[1], Indira Pradasadam[1-2], Yi-Chin Toh[1-4]

[1]School of Mechanical, Medical and Process Engineering, Queensland University of Technology, Brisbane, Australia

[2]QUT Centre for Biomedical Technologies, Queensland University of Technology, Brisbane, Australia

[3]Max Planck Queensland Centre (MPQC) for the Materials Science of Extracellular Matrices, Queensland University of Technology, Brisbane, Australia

[4]Centre for Microbiome Research, Queensland University of Technology, Brisbane, Australia

### Supplementary Information

#### SI Table 1: Antibody list

| Antibody | Host | Vendor | Catalogue number |
|---|---|---|---|
| HIF-1α | Mouse | ThermoFisher Scientific, Australia | MA1-516 |
| COL1A | Rabbit | Abcam, Australia | AB138492 |
| COL2A | Mouse | Abcam, Australia | AB185430 |

#### SI Table 2: Secondary Antibody list

| Antibody host | Vendor | Catalogue number | Excitation / emission (nm) |
|---|---|---|---|
| Donkey anti-mouse | ThermoFisher Scientific | A32773 | 555/600 |
| Donkey anti-rabbit | ThermoFisher Scientific | A32790 | 488/520 |

#### SI Table 3: Primer sequence from IDT Australia

| Name | Sequence (5'-3') | |
|---|---|---|
| | Forward | Reverse |
| *18s* | TTC GGA ACT GAG GCC ATG AT | CGA ACG TCC GAC TTC GTT C |
| *COL1A1* | CCC TGG AAA GAA TGG AGA TGA T | ACC ATC CAA ACC ACT GAA ACC T |
| *COL1A2* | GCA ACA TGC CAA TCT TTA CAA GAG | CCA TCA TAC TGA GCA GCA AAG TTC |
| *COL2A1* | TGG ACG ATC AGG CGA AAC C | GCT GCG GAT GCT CTC AAT CT |
| *AGAN* | GAG ACA CCA ACG AGA CCT ATG ATG | GCA CTC ATT GGC TGC TTC CT |
| *ADAMTS5* | TGG CTC ACG AAA TCG GAC ATT | TGC ATT TGG ACC AGG GCT TA |

| | | |
|---|---|---|
| *MMP13* | TCA CCA ATT CCT GGG AAG TCT | TCA GGA AAC CAG GTC TGG AG |
| *ALP* | TCT TCA CAT TTG GTG GAT AC | ATG GAG ACA TTC TCT CGT TC |
| *SSP1* | TCA CCA GTC TGA TGA GTC TCA CCA TTC | TAG CAT CAG GGT ACT GGA TGT CAG GTC |

**SI Table 4: Curated list of genes with reported regulatory functions in cartilage metabolism according to reported literature [42-45]**

| **Catabolic activity** | **Anabolic activity** |
|---|---|
| IL1B | IL13RA2 |
| IL17RC.21 | IL10RA |
| IL17RC.22 | IL10RA.2 |
| IL18 | TGFB3 |
| LIF.2 | TGFBR2 |
| LIF.4 | TGFBR2.2 |
| IL6 | IGF2.3 |
| IL8 | IGFBP5 |
| MMP10 | IGF1 |
| MMP1 | IGFBP4 |
| MMP9 | IGF2 |
| ADAMTS1 | COL6A1 |
| TNFAIP8 | COL2A1.3 |
| TNFAIP3.3 | COL3A1 |
| TNFAIP2.2 | COL4A2 |
| TNFAIP3.5 | MXRA8 |
| IL17RB | FGFR2.4 |
| IL17D.3 | FGFR2.8 |
| IL17D.2 | BMP6 |
| IL18BP.4 | TGFB1 |
| LIF | IGFBP1 |
| MMP3 | IGF2BP1.2 |

| | |
|---|---|
| MMP11 | IGF1.4 |
| MMP12 | SOX9 |
| MMP15 | COL16A1 |
| MMP13 | MAT2A |
| ADAMTS16 | COL1A1 |
| TNFAIP6 | COL1A2 |
| IL17D | IL8 |
| SMAD1.4 | BMP2K.3 |
| SMAD3.3 | TGFBI |
| SMAD9.2 | IGF1.5 |
| SMAD9 | NFKBID.2 |
| | BMP6 |

**Supplementary Figures**

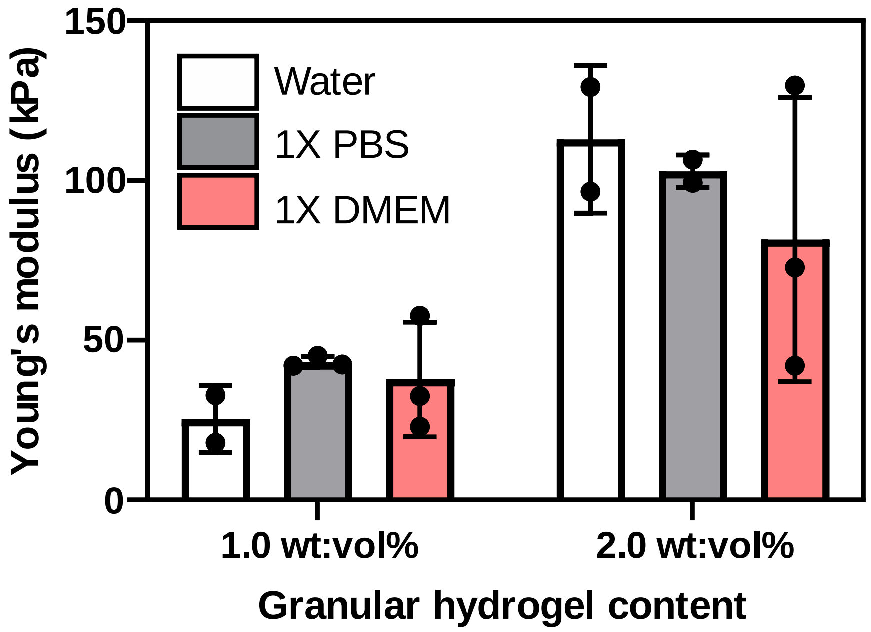


**SI Figure 1:** Characterization of granular hydrogel stiffness prepared with DI water, 1X phosphate buffer saline (PBS) and 1X DMEM culture medium. The hydrogel stiffness can be tuned with the weight % of the polyacrylate content as well as the liquid content. Data are average ± S.D. of 3 independent samples.

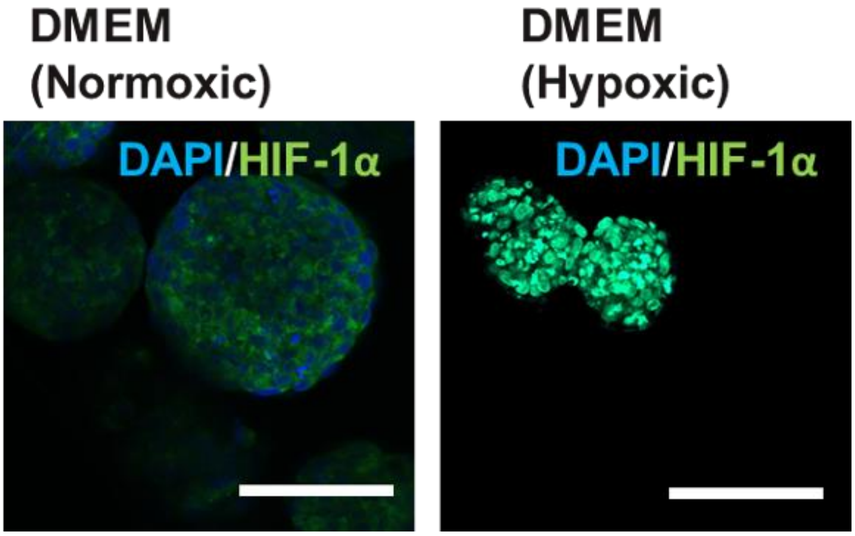


**SI Figure 2:** DAPI and HIF-1α staining of human primary chondrocytes culture in liquid low glucose DMEM in regular incubator (normoxic) and hypoxic incubator (hypoxic) after 4-day culture. Scale = 100 µm.

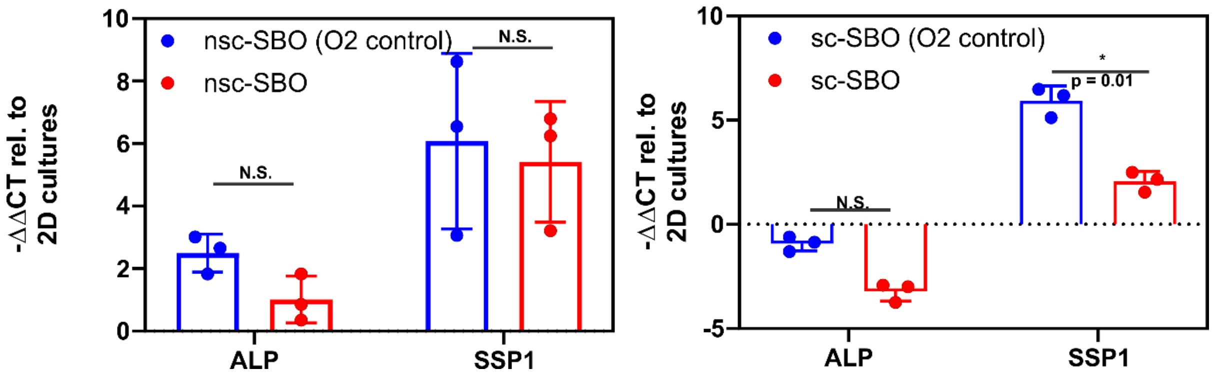


**SI Figure 3:** Gene expressions of alkaline phosphatase (*ALP*) and osteopontin (*SSP1*) in non-sclerotic osteoblasts (nsc-SBO) and sclerotic osteoblasts (sc-SBO) after 4 days of co-culture with primary chondrocytes under differential (2%-20%) and uniform (20%) oxygen levels. *ALP* and *SSP1* expressions in nsc-SBO and *ALP* expression in sc-SBO were not significantly different regardless of whether they were patterned next to chondrocytes in hypoxic or normoxic hydrogels. *SSP1* expression was significantly higher in sc-SBO co-cultured with chondrocytes in hypoxic hydrogels. All gene expressions were normalized to expression levels of 2D cultures of osteoblasts at 20% oxygen. Data are individual experimental of replicates of n = 3 with S.D,

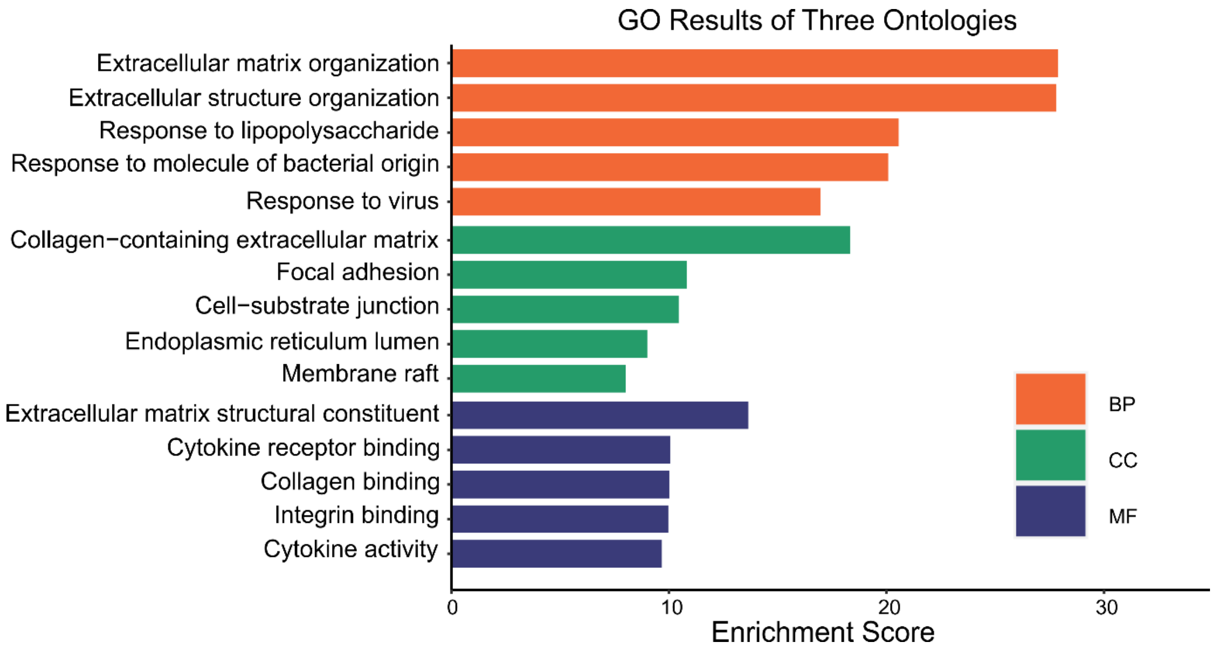


**SI Figure 4:** Gene ontology (GO) analysis of the RNA-sequencing data for biological processes (BP), cellular components (CC) and molecular functions (MF). The chondrocytes were co-cultured under hypoxia
